# DeepImageJ: A user-friendly environment to run deep learning models in ImageJ

**DOI:** 10.1101/799270

**Authors:** Estibaliz Gómez-de-Mariscal, Carlos García-López-de-Haro, Wei Ouyang, Laurène Donati, Emma Lundberg, Michael Unser, Arrate Muñoz-Barrutia, Daniel Sage

## Abstract

DeepImageJ is a user-friendly solution that enables the generic use of pre-trained deep learning (DL) models for biomedical image analysis in ImageJ. The deepImageJ environment gives access to the largest bioimage repository of pre-trained DL models (BioImage Model Zoo). Hence, non-experts can easily perform common image processing tasks in life-science research with DL-based tools including pixel and object classification, instance segmentation, denoising or virtual staining. DeepImageJ is compatible with existing state-of-the-art solutions and it is equipped with utility tools for developers to include new models. Very recently, several train ing frameworks have adopted the deepImageJ format to deploy their work in one of the most used software in the field (ImageJ). Beyond its direct use, we expect deepImageJ to contribute to the broader dissemination and reuse of DL models in life-sciences applications and bioimage informatics.

---

Deep learning (DL) models have a profound impact on a wide range of imaging applications, including life-sciences [1, 2]. Unfortunately, their accessibility is often riddled with technical challenges for the non-expert user. Since most DL methods are available as source code, running them requires setting up sophisticated software and hardware environment. The increasing use of image analysis workflows in biomedical research [1] and the willingness to disseminate trained DL models have pushed computer scientists to design more user-friendly solutions [3, 4, 5]. Currently, there exists an increasing number of active developer teams addressing this problem with different solutions: the CSBDeep team [6]^1^, the Ozcan Research Group^2^, DeepClass4Bio [7]^3^, Ilastik [8]^4^, ImJoy [9]^5^, ZeroCostDL4Mic [10]^6^, YAPIC [11] and DeepTrack [12]. The CSBDeep team distributes their DL workflows via an ImageJ [3, 13] toolbox [6] which lets non-expert users perform a variety of microscopy image analysis using trained DL models. Through their plugin, it is possible to train denoising models in a local machine without any previous programming skills. The StarDist plugin [14] makes the most powerful tool for cell nuclei detection and segmentation in microscopy images accessible in ImageJ. Similarly, the Ozcan Research team has often made its trained models available in ImageJ [15]. DeepClass4Bio is an API to use image classification tasks in ImageJ and Icy^7^ using trained DL models. The Ilastik team has an early release of a neural network classification workflow equipped with both inference and training functionalities. ImJoy [9] is a generic computational platform to deploy advanced data anlysis applications and make them accessible to non-expert users. It is particularly suited for building and sharing interactive web interfaces for DL-based image analysis. The ImJoy ecosystem allows the integration of a variety of open-source software such as ImageJ, so it can be used without previous installation. ZeroCostDL4Mic [10] utilizes the free cloud GPU resources provided by Google Colaboratory^8^ and provides extensive documentation in a browser-based notebook interface, allowing non-experts to train DL models such as a generic segmentation model (*e*.*g*., the well-known U-Net [16]) or the super-resolution microscopy model (*e*.*g*., DeepSTORM [17]). YAPIC is a Python library to train a U-Net on pixel classification and make predictions by writing few plain command lines. DeepTrack combines browser user interfaces to train different models in a non-coding fashion, together with a set of Python Notebooks that support the easy training and use of their models.

The previously mentioned tools have started to boost the use of DL solutions for biomedical analysis tasks. However, a user-friendly tool to disseminate trained models for image processing in a non-coding fashion and with a unified format is still missing [1, 18]. We present *deepImageJ*^9^, an open-source environment for ImageJ which is the *de-facto* standard image processing software in life-sciences [3]. The open-source package ImageJ gives biologists access to a wide variety of user-friendly image analysis tools through third-party plugins and macros. It contains most of the standard bioimage analysis methods and is continuously updated with the most recent techniques. The current integration of deepImageJ further contributes to the ImageJ ecosystem. Since the first release of deepImageJ, the framework has been widely used by developers to share their work with collaborators in the life-sciences domain or to provide an easy way to test a DL solution for their biological imaging application (see Figure 1). DeepImageJ runs a variety of third-party models from the main DL libraries that are powering the DL framework (TensorFlow^10^ and PyTorch^11^). Installing deepImageJ is straightforward compared to that of common Python environments thanks to the one-click installation facilitated by the ImageJ updates manager: deepImageJ can be installed through ImageJ’s update site. DeepImageJ is designed as a standard ImageJ plugin with the technicalities hidden behind the user-friendly interface. We provide detailed instructions to facilitate its configuration. [Video supplementary material]. Similarly to any other ImageJ plugin, deepImageJ can run in any local machine, independently of its Operating System (OS). An advantage compared to cloud-based tools such as Google Colaboratory, is that deepImageJ runs locally without the need of uploading data, thus providing better privacy protection.

**Figure 1:**
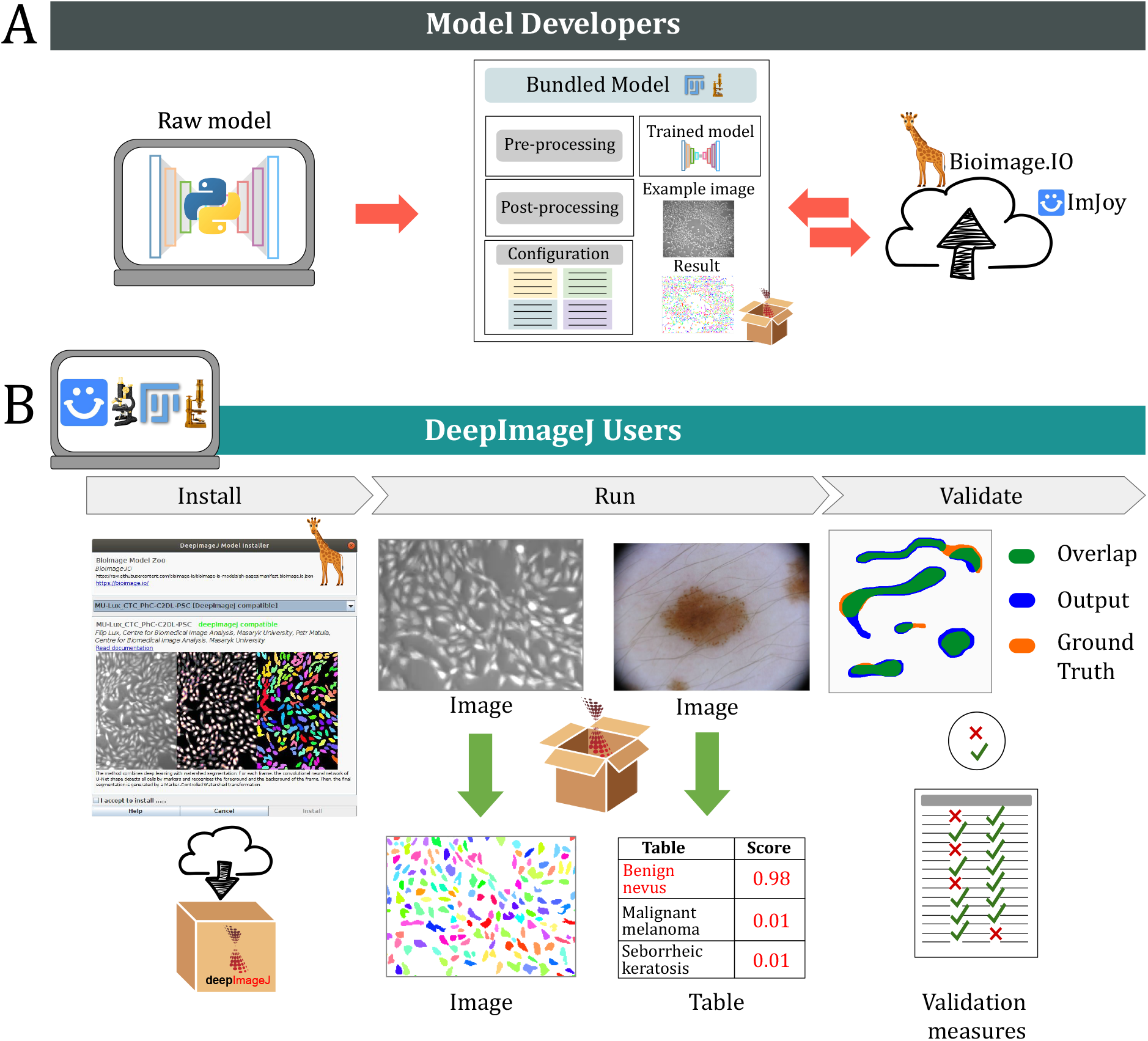
DeepImageJ environment and scope. DeepImageJ targets model developers and non-expert biomedical image analysts. A) Developers train a Deep Learning (DL) model usually in Python and bundle it using the `DeepImageJ_Build_Bundled Model` plugin in ImageJ. The bundled model follows the BioImage Model Zoo format: it contains the trained model architecture and weights, pre- and post-processing routines, testimonial images for reproducibility purposes and a specifications file. The latter gathers all the technicalities that allow cross-compatibility among the BioImage Model Zoo consumer software, deepImageJ among others. The bundled model package can be disseminated through the public model repository in the cloud synchronized with the BioImage Model Zoo, or directly sending it to the final user through a private communication channel. B) Life-scientists can run a deepImageJ model online using ImJoy, or install it locally. The model can be used as any regular ImageJ plugin to analyze images. The deepImageJ model package allows the automatic processing of any input image and is compatible with outputs of different formats and dimensions such as images or tables. The `DeepImageJ_Validate` plugin provides a set of evaluation measures to compare the resulting image with a ground-truth image provided by the user.

DeepImageJ operates with several types of models for image processing tasks such as image-to-vector (*e*.*g*. image classification), image-to-image (*e*.*g*. image segmentation, deconvolution, virtual labeling, super-resolution) or pyramidal feature pooling networks (*e*.*g*. region proposal networks for object detection, panoptic segmentation) (see Figure 2). This is possible due to the technical specification of a DL model that is independent of the bioimage analysis task or the model architecture. Succinctly, the deepImageJ format is as general as possible so that the use of different models does not rely on any technical configuration or work, i.e. utilizing one model or another, makes no difference to the user. The latter goes in hand with the recent BioImage Model Zoo ^14^ initiative which is meant to unify the effort of the biomedical image analysis community to make accessible, open, and usable the trained DL model through different consumers. As part of the initiative, we are building an open-source repository to deliver trained DL models, Python notebooks, and biomedical image datasets in a standard manner, resulting in a combined effort for the democratization of DL in the biomedical image analysis field. The deepImageJ format has migrated to the specifications of the BioImage Model Zoo, allowing interoperability between the community partners software, namely, Ilastik, CSBDeep, deepImageJ, ImJoy and ZeroCostDL4Mic. Likewise, the deepImageJ model repository is synchronized with the BioImage Model Zoo. The first selection of deepImageJ-compatible trained models is available in the repository, from where the user can easily download and apply in ImageJ the model of its choice.

**Figure 2:**
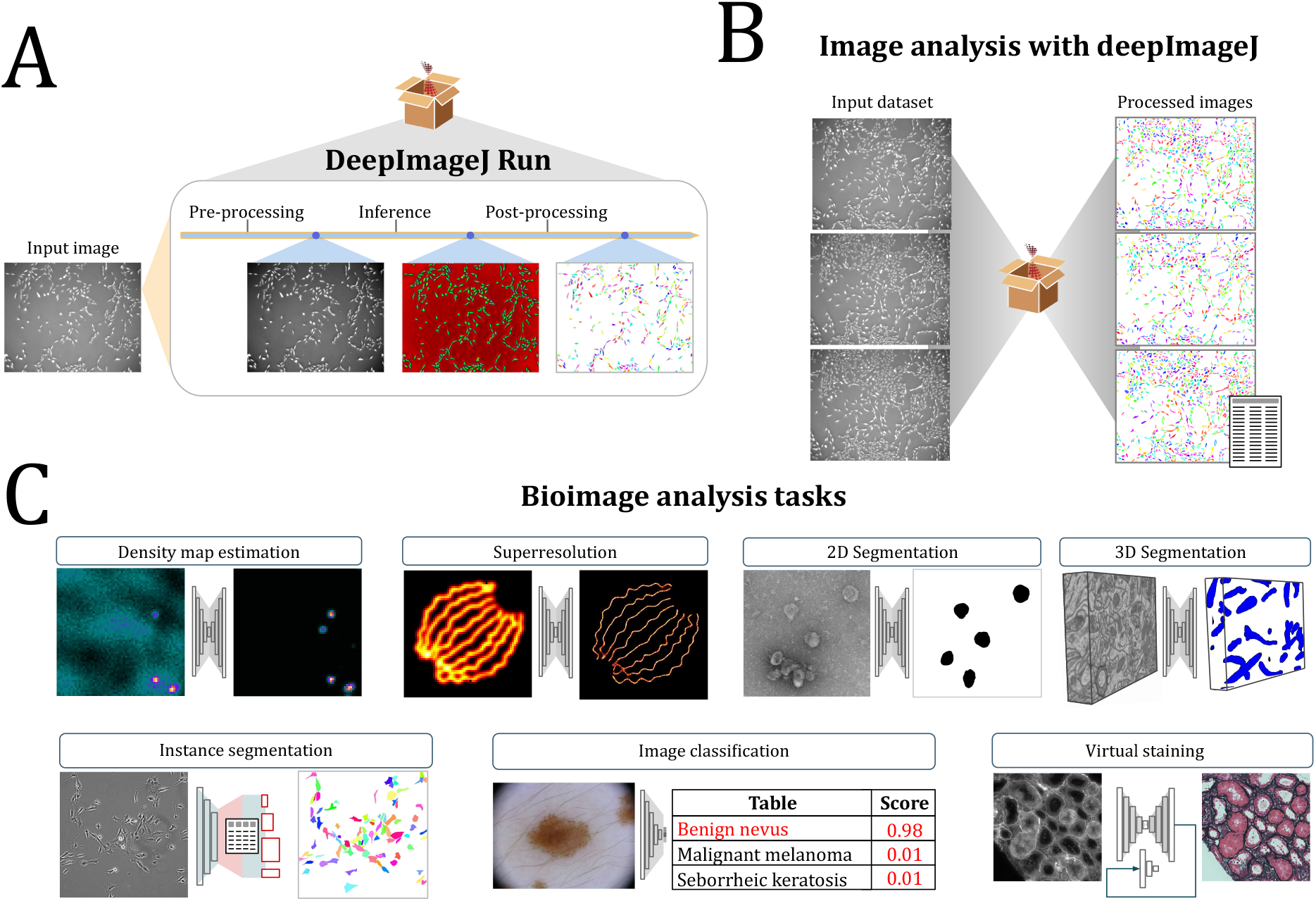
Functionalities of deepImageJ: A) `DeepImageJ_Run` plugin processes an input image with a locally installed deepImageJ compatible trained model. It automatically carries out the pre-processing, inference and post-processing steps required and written previously by a developer; B) The `DeepImageJ_Run` plugin can be called from ImageJ macros, so locally stored image datasets can be automatically processed. Furthermore, the use of deepImageJ trained models can be integrated into extended bioimage analysis pipelines; C) DeepImageJ targets those deep learning models that have an image as input and it is compatible with outputs of different formats and dimensions. Therefore, it is suitable for tasks such as density estimation^12^, superresolution [17], 2D and 3D segmentation [19, 16], instance segmentation [20], image classification^13^ or virtual staining [15].

The main plugin of the deepImageJ toolbox, `DeepImageJ_Run`, allows the execution of DL models for an image processing task in a few clicks without DL expertise or programming skills. During this process, users have access to a description and the complete documentation of the trained model. If a GPU is locally available and set up following the guidelines in deepImageJ’s website, the plugin will automatically connect with it. Image-to-image processing tasks such as segmentation, denoising, deconvolution or super-resolution, need a tiling strategy when using DL models (see Online Methods). The plugin guides the user through this step by recommending a specific tile size whenever possible. In addition, the plugin ensures the consumption of pre-processing, inference and post-processing routines as detailed by the model developer (see Figure 2). While the process is automatically executed, the plugin informs the user about each performed step. This provides flexibility to users who want to test different model configurations and understand more in-depth the entire process. Moreover, `DeepImageJ_Run` can be called from the popular scripting ImageJ macro, which allows the use of the models in larger image processing workflows. This feature facilitates a variety of setups such as the automatic processing of a locally stored set of raw microscopy images^15^. See Figure 2.

The deepImageJ toolbox is complemented by three companion plugins for model bundling, model installation and validation of the results:

- `DeepImageJ_Build_BundledModel` guides model developers to bundle their trained model and provide all the necessary information (meta-data) for its easy use in ImageJ (see Online Methods). The plugin creates the package required to upload the model to the online repository (see Figure 1). Alternatively, model developers can use the `pydeepimagej`^16^ library to create the bundled models directly from the source code in Python.
- `DeepImageJ_Install_Model` allows to access the BioImage Model Zoo repository to install a model stored in the cloud. (See Figure 1).
- `DeepImageJ_Validate` tool gives access to a variety of perceptual and segmentation validation measures that guide the user through an evaluation of the obtained result. See Figure 1.

To facilitate deepImageJ model testing, we ported `DeepImageJ_Run` plugin to Javascript such that it can run in a web browser via ImageJ.JS (https://ij.imjoy.io). This enables the integration with ImJoy, which allows testing models in the BioImage Model Zoo website without downloading any model or installing the plugin locally. This is especially helpful for users to compare and select models based on their data, making the dissemination of DL models more effective.

When running DL models, it is crucial to pre-process the input image so the model is fed with an image that has the same precise features as the ones employed in the training process (*e*.*g*., data normalization pre-processing). To handle this, deepImageJ gives the user the flexibility to run pre- or post-processing routine written in an ImageJ scripting language: ImageJ macro or Java plugins. The latter is the main bridge between complex architectures (for example, the Mask-RCNN employed in Usiigaci [20]) and our non-code solution in ImageJ. (See Figure 2).

Although we made every effort to lay solid foundations to use the deepImageJ toolbox, the correctness of a DL model’s output ultimately depends on its appropriate usage. Hence, it is critical that the user pays close attention to the information given by the DL developers before running a model, and that all the results obtained are thoroughly inspected. For this, users can take advantage of `DeepImageJ_Validate` to assess the accuracy of the results whenever ground truth data is available. Unlike in other computer vision niches, most DL models for microscopy image processing still lack the ability to generalize across datasets [18]. While we are confident about the future developments to get general and data-cross-compatible models, it is currently recommended to fine-tune the DL models. Such (re)training and evaluation of a model is only possible when there is ground truth data, proper infrastructure (software and hardware) and most often, knowledge about machine learning. Because model training and fine-tuning are out of the scope of deepImageJ, we would like to highlight already existing user-friendly tools like Ilastik, ImJoy, ZeroCostDL4Mic or YAPIC which can perform such tasks. We work on the cross-compatibility of some of these tools to provide combined solutions. For example, using ZeroCostDL4Mic, the U-Net [16]) and DeepSTORM [17] models, which are well-established DL models for biomedical image processing, can be trained, automatically exported and used with deepImageJ. Yet we can find another example of linking deepImageJ with DL model training to provide an end-to-end workflow of cell morphology analysis^17^. The YAPIC team has integrated a new feature in the software to export any of their models into the deepImageJ format. We encourage all users with a potential need of fine-tuning their models, to integrate these combinations in their pipelines. Moreover, such combinations are becoming popular in bioimage analysis as they exploit the computational capacity of Python to easily (re)train DL models and the flexibility given in tools such as ImageJ to concatenate different image processing steps.

The use of trained DL models without a deeper understanding of the method could potentially become the source of unreliable results. Aware of the latter, we remark the very recent effort made by the community to define and recommend good *praxis* in bioimage analysis, and more specifically, to train life-scientists on both the promises and risks of using this new technique. We trust in deepImageJ’s potential as a user-friendly tool to bridge the current gap between computer vision and biomedical image analysis. The deepImageJ environment is highly suitable for educational activities due to its easy installation and model execution, and the flexibility achieved thanks to access to all the ImageJ resources.

The experience gathered through the development of deepImageJ has seeded the scientific collaborations among different actors involved in the bioimage analysis field and set a roadmap for future developments, with key aspects being interoperability and smart guidance towards automated machine learning (AutoML) [1]. Because DL models consist of numerous layers and operations, using them to process images requires considerable memory consumption. DeepImageJ integrates a smart feature that allows adjusting the patch size when possible, to smaller image size and complete the tiling strategy automatically without further changes in the model specifications. Nonetheless, further improvements can still be done to design an intelligent memory management strategy for running the DL models, such as adjust the input patch shape automatically according to the user’s resources. DeepimageJ, for example, provides relevant information for the user to know how much memory will the model execution consume, and therefore, to determine if the model can be used in a certain machine. However, it will not set up the model execution configuration according to the memory capacity. A straightforward example is the smart configuration of the tiling strategy considering the capacity of the user’s machine. Yet another challenge is the development of personalized guides based on the user’s data to choose the best-suited model and proper fine tuning strategies. So it is that AutoML could turn into the new hot-topic in bioimage analysis.

Notwithstanding the current tendency to build general models for image processing tasks [18], there is a pressing need to release user-friendly and model-adjustable environments to run DL models [1, 18, 4]. The latter goes in hand with a higher-level target of reducing human interaction in the routinely performed bioimage analysis tasks. That said, deepImageJ is a contribution to the field for the accessibility of DL in bioimage analysis and the dissemination of DL models in a standardized manner. The deepImageJ environment facilitates the work exchange between developers and end-users, which the ongoing integration of deepImageJ in the BioImage Model Zoo seeks to reinforce. Therefore, we expect deepImageJ to enlighten the work of life-sciences researchers and to become yet another standard open-source tool for the dissemination of DL image processing models.

## Data availability statement

The web page: https://deepimagej.github.io/deepimagej/ provides free access to the plugin, along with the bundled models and user guide for image processing.

## Acknowledgments

We would like to thank Alexandre Levy from École polytechnique fédérale de Lausanne (EPFL) for his contribution to the validation tool of deepImageJ. Pedro M. Gordaliza, Ignacio Arganda-Carreras (tested the beta-versions), Daniel Felipe González-Obando (tested the beta-versions), Curtis Rueden, Sébastien Tosi, Thomas Pengo (tested the beta-versions), Ricardo Henriques (ZeroCostDL4Mic), Romain F. Laine (ZeroCostDL4Mic), Guillaume Jacquemet (ZeroCostDL4Mic), Daniel Krentzel (ZeroCostDL4Mic), and Christoph Möhl (YAPIC) for the fruitful discussions and enriching feedback about the deepImageJ project. We would also like to thank NEUBIAS for supporting the project, the NEUBIAS symposium and NEUBIAS Academy@Home. Pejman Rasti and Steffen Bollmann for including deepImageJ in their tutorials. We thank the reviewers of Nature Methods for their constructive comments. We would like to specially mention all the contributors and community partners at the BioImage Model Zoo for the time they have spent to get a cross-compatible model format.

We acknowledge the support of Ministerio de Ciencia, Innovación y Universidades, Agencia Estatal de Investigación, under Grants TEC2016-78052-R and PID2019-109820RB-I00, MINECO/FEDER, UE, co-financed by European Regional Development Fund (ERDF), “A way of making Europe” (EGM, CGLH, AMB), and a 2017 Leonardo Grant for Researchers and Cultural Creators, BBVA Foundation (AMB). This work was also supported by the EPFL Center for Imaging (CGLH, LD, DS, MU). We would like to thank the Science for Life Laboratory, Erling-Persson Foundation and Knut and Alice Wallenberg foundation (2018.0172)(Wo, EL). We thank the program “Short Term Scientific Missions” of NEUBIAS (network of European bioimage analysts)(EGM, CGLH). We also want to acknowledge the support of NVIDIA Corporation with the donation of the Titan X (Pascal) GPU card used for this research (AMB).

## Author contributions

E.G.M. and C.G.L.H. contributed to the design of the experimental framework, reviewed, trained and exported existing image processing methods. C.G.L.H. and D.S. developed and implemented the toolbox and worked on the supporting documentation with input from the rest of the authors. C.G.L.H. and W.O. built the connection between the toolbox and ImJoy. E.G.M. wrote the code lines of the supplementary Python notebooks, Python library and ImageJ macros. E.G.M. and W.O. worked on the synchronization with the BioImage Model Zoo. E.G.M., W.O. and L.D. wrote the manuscript with help from E.L., M.U., A.M.B. and D. S. E.G.M., and D.S. created the website of deepImageJ. M.U., A.M.B., and D.S. initiated the project. A.M.B. and D.S. supervised the project. All the authors contributed to the conception of the study, the design of the experimental framework and took part in the literature review. All authors revised the manuscript.

## ONLINE METHODS

DEEPIMAGEJ: A USER-FRIENDLY ENVIRONMENT TO RUN DEEP LEARNING MODELS IN IMAGEJ

In the following lines technical descriptions and software requirements are provided. The ImageJ user guide for this version of the plugin (deepImageJ 2.1.0) is given in the Supplementary Material. As deepImageJ is an on-going project, we strongly recommend to check the documentation at deepImageJ’s website^1^ and the plugin’s Wiki in GitHub^2^.

## 1 Software/Network compatibility

DeepImageJ is compatible with both Fiji [1], ImageJ [2] and ImageJ2 [3]. It is self-sufficient on any operating system: MacOS, Linux and Windows and on 64-bit operating systems (32-bit operating systems are not supported). It supports TensorFlow models until version 1.15, and PyTorch 1.6. Keras version 2 or lower is also supported as long as the models are compatible with Tensorflow version 1.15 or lower. Same as CSBDeep [4], deepImageJ uses a TensorFlow Java API manager to ensure TensorFlow version compatibiltiy. The latter can be upgraded in ImageJ2 through the ImageJ-TensorFlow manager^3^, developed by Curtis Rueden and Deborah Schmidt. The Deep Java Library^4^ ensures the compatibility with PyTorch. Nonetheless, those users with Windows operating system need to install Visual Studio^5^ for the deployment of this library in ImageJ/Fiji.

The Java libraries used to load TensorFlow and PyTorch models point to the same source code as the respective Python packages. This implies that regardless the code (Python or Java), the same results and execution times are ensured.

## 2 DeepImageJ bundled models

DeepImageJ Run processes folders (models) that contain the files described in Table 1. Those files are usually written by the author of the model, which makes the bundled model self-sufficient. Their content is described in the next paragraph. Further details about loading bundled models in deepImageJ are given in the Supplementary Material.

**Table 1:** List of the required files to run a model in deepImageJ.

| File name | Content |
| --- | --- |
| saved_model.pb<br>variables<br>OR<br>pytorch_model.pt | TensorFlow model in Protocol Buffer format (saved bundled model).<br>PyTorch model stored in TorchScript format. |
| file(.ijm / .jar / .class) | ImageJ macros<sup>8</sup> or Java code written by the model's author for the pre- and post-processing. |
| exampleImage.tif | Example of input the image. |
| result(.tif / .csv) | Output of the model after the post-processing |
| model.yaml | Bioimage Model Zoo configuration specifications: Contains details about the related publication and technical characteristics of the model. |

Any Deep Learning (DL) model is determined by a graph (the architecture of the network) and its weights (specific values for all the parameters in the network obtained after training). The TensorFlow’s Java API is only compatible with the SavedModel format which is obtained using an in-house Python routine^6^. Namely, the deepImageJ models are defined by a Protocol Buffer format file (`called saved_model.pb`) that contains the architecture of the model and a series of text files storing the weights that are kept in a folder called `variables`. PyTorch models should be exported in TorchScript format (`pytorch_model.pt`) which contains both the architecture and the weights compressed in a single file. ImageJ macros (`.ijm`) or compiled Java code (`.jar / .class`) are optional pre- and post-processing steps. The pre-processing routine transforms the image into a specific input type for which the model was trained. Typical pre-processing operations are normalization of the pixel intensity values, change of the bit depth and image resizing. The post-processing routine curates the output of the network. Optional post-processing operators are for example, thresholding, resizing or extraction of objects features. At least two files, an input image (`exampleImage.tiff`) and an output result (`.tif / .csv`) are also stored in each of the bundled model folder to facilitate model testing.

The configuration file (model.yaml) has descriptive information about the model and it is synchronized with the Bioimage Model Zoo configuration specifications^7^:

- General information: Name of the author(s), title, description, reference to the publication or GitHub repository, license and framework.
- Technical characteristics of the model: Input and output specifications (dimensions shape, size and axis order), pre- and post-processing information and model format.
- DeepImageJ specific information: Example image name and pixel size, TensorFlow signature name, pre- and post-processing filenames, minimum amount of memory required to process the example input image, estimated execution time on the PC.

### 2.1 Interpretation of the configuration specification (model.yaml)

The configuration specifications file has a differentiated section under the field name config in which each software can include its custom specifications without altering the common format that is compatible with the community partners at the Bioimage Model Zoo. For deepImagej, the config field follows the next schema

**Listing 1:**
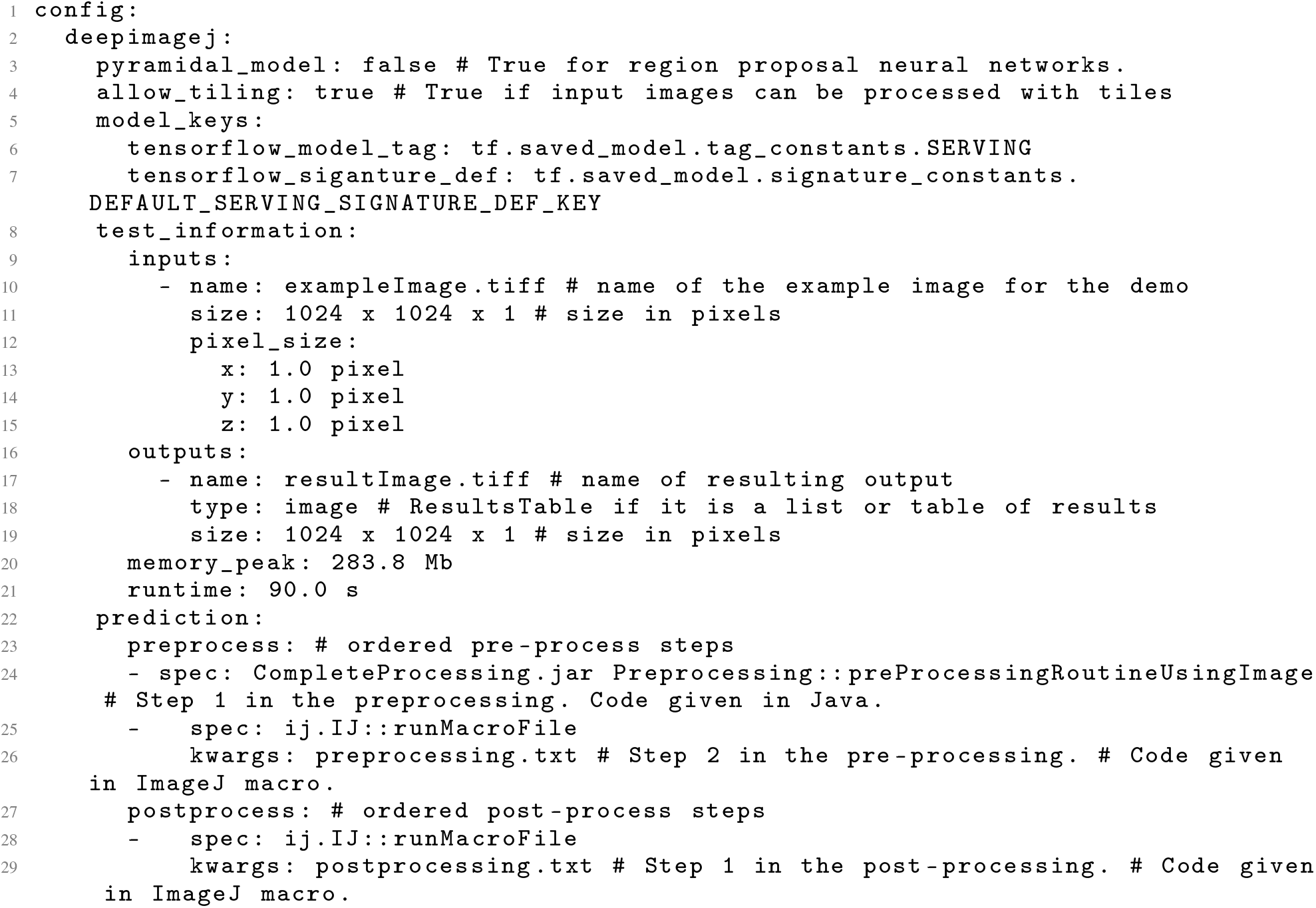
DeepImageJ specific configuration in model.yaml

## 3 Input and output size calculations: Tiling strategy

The model developer needs to specify the following information when uploading their Convolutional Neural Network (CNN) model:

*Q, I*: Whether the input size of the model (*Q*) is predetermined or not. If it is predetermined, *Q* needs to be provided, and it will be compared with the size of the image to process (*I*). *Q* corresponds to the field called `shape in the model.yaml`.

*m, s*: If the network has an auto-encoder architecture, the size of each dimension of the input image, has to be multiple of a minimum size *s* defined as *s* = *p*^*d*^ where *d* is the number of `poolings` (down-sampling operations) and *p* their size. Additionally, the input should have a minimum size *m* so the shape of the tensor that will enter the model satisfies the equation

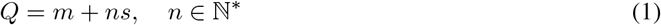

*m* and *s* correspond to the fields called min and step respectively in the `model.yaml`.

*P*: To preserve the input size at the output, convolutions are usually calculated using zero padding boundary conditions (Figure 1 is an illustration of the 2D case). Namely, additional void pixel values are added along the borders of the image. Hence, the size (per dimension) of the valid domain of the output is given by *R* = *Q* –2*P* with *Q*, the model input size and *P*, the size of the network padding, sometimes denoted as halo. The size of the padding is equivalent to the receptive field of one pixel in the CNN. For a symmetrical encoder-decoder architecture, namely, the same number of down and up-samplings. It can be computed as

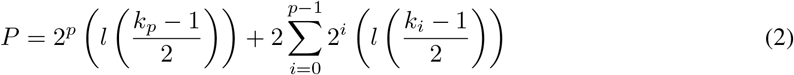

where *k*_*i*_ is the kernel size for each convolutional layer, *p* is the number of poolings, and *l* is the number of convolutional layers at each level of the encoder-decoder. Usually, *k*_*i*_ is an odd number. If the kernel is not square, then *P* and *R* have different values on each dimension. *P* is given in the field halo of the model.yaml.

To handle input images with a large size, deepImageJ follows a common strategy called *tiling*:

- If the network has not a predetermined input size (*Q*), the algorithm calculates what is the smallest size *t* that satisfies Equation 1 and is still larger or equal to the size of the input image *I* and the total padding 2*P* :

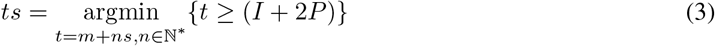

Then, the image is augmented by mirroring along the borders up to a size *t* per dimension, and it is processed. Finally the output is cropped to the initial size *I*. See Figure 1 for an illustration.
- If the network input size (*Q*) is predetermined, then the algorithm compares the size of the image (*I*) with it taking into account the padding (*P*). If it is smaller (*I* + 2*P* ≤*Q*), then the image is augmented by mirroring until the desired size *Q* is reached. If the opposite is true (*I* + 2*P > Q*), the optimal number of tiles to process (*N*) is calculated as follows:

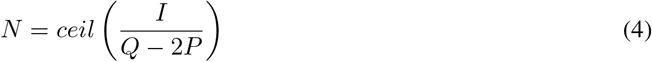

where the function *ceil*(*x*) outputs the smallest integer number that is equal or larger than *x*. Note that *N* can vary on each dimension. Then, the image will be covered by patches of size

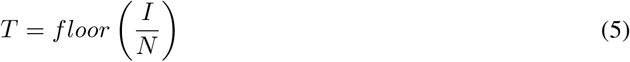

where the function *floor*(*x*) outputs the largest integer number that is equal or smaller than *x* and *T*≤ (*Q*–2*P*). From each processed patch of size *Q*, a patch of size *T* is cropped and placed accordingly to reconstruct a valid output (tiling strategy). The patches along the borders are filled by mirroring as shown in Figure 1. As the quotient in Equation 5 may not be an entire number, the last patch on each dimension has exactly size *I* – (*N* – 1)*T*.

**Figure 1:**
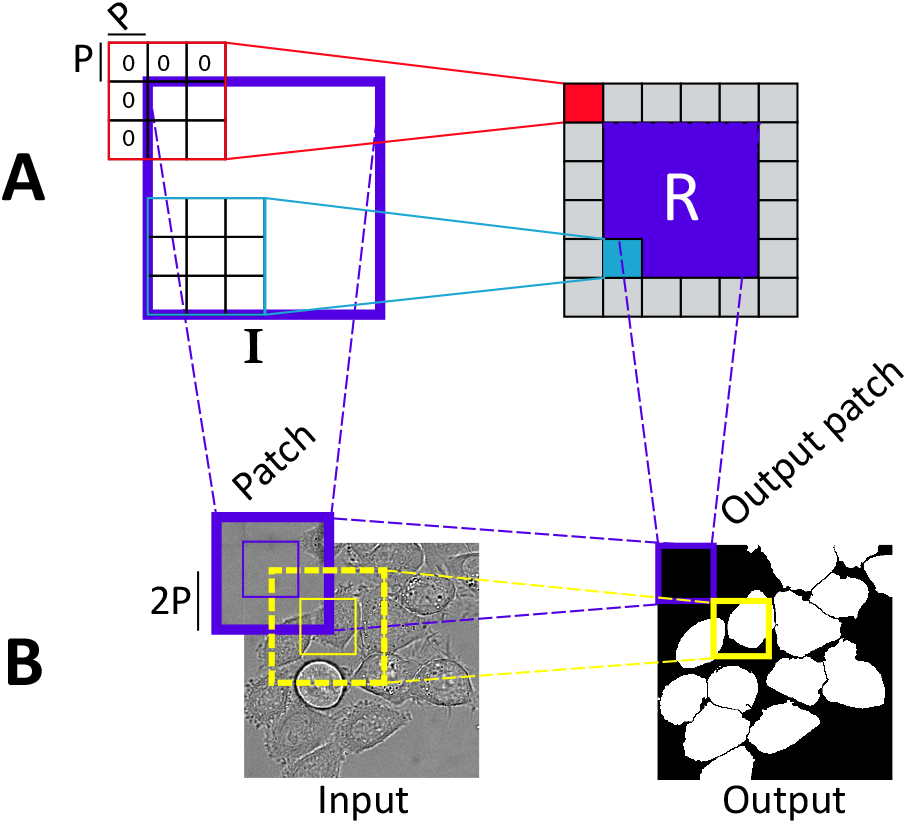
Tiling strategy with convolutions. **Panel A:** Convolution of size 3 *×* 3 pixels. The receptive field of a convolutional layer, *R*, is smaller than the input image and it is determined by the size of the padding (red operation) used to get an output image of the same size as the input image. **Panel B:** Image processing using tiles (patches). The overlap 2*P* is calculated to avoid artifacts in the reconstruction of the output.

Both the input size of the network (*Q*) and the padding of the CNN (*P*) are critical parameters for good results and they are directly related to the time spent by the plugin to process one image. Large input images and deeper networks that have a larger receptive field, imply longer computations.

The current deepImageJ version also handles inputs and outputs of different sizes. As for example, the case of models for super-resolution. To determine the shape of the output image given a certain input image, we do

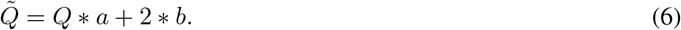

where *a* and *b* are the `scale` and `offset` parameters respectively given in the `model.yaml` file, *Q* the shape of the input image, and 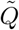 the shape of the output image. Likewise, in the model.yaml file, the reference image for *Q* needs to be indicated. The latter is given in the field `reference_input`.

## 4 Examples of a complete image analysis pipeline

In the following lines, we point the reader to different workflows for image analysis using DL models and deepImageJ plugin.

User-oriented pipelines to train a DeepSTORM model for Single Molecule Localization Microscopy (SMLM) or object segmentation using 2D and 3D U-Nets are provided by ZeroCostDL4Mic^9^ [5]. These notebooks guide the user through all the classical steps in DL (data preparation, data augmentation, training, evaluation and inference of new images). Additionally, they provide a specific section to export the trained model into the deepImageJ format. In [6], Gómez de Mariscal et al. describe how to build from scratch a full pipeline with Python programming for cell segmentation in phase contrast microscopy images. The pipeline uses the Google cloud services to train the model and then it shows how to work with that model locally using deepImageJ, so it avoids uploading the analyzed data to Google Drive.

Yet another example is as follows:

We use a toy example in which a U-Net [7] is trained to segment glioblastoma cells on 2D phase contrast microscopy images. The entire code was written in Python using the Keras library^10^ and it was run in the Google Colaboratory environment, which supplies a free Tesla K80 GPU service. The code is freely distributed^11^.

**Figure 2:**
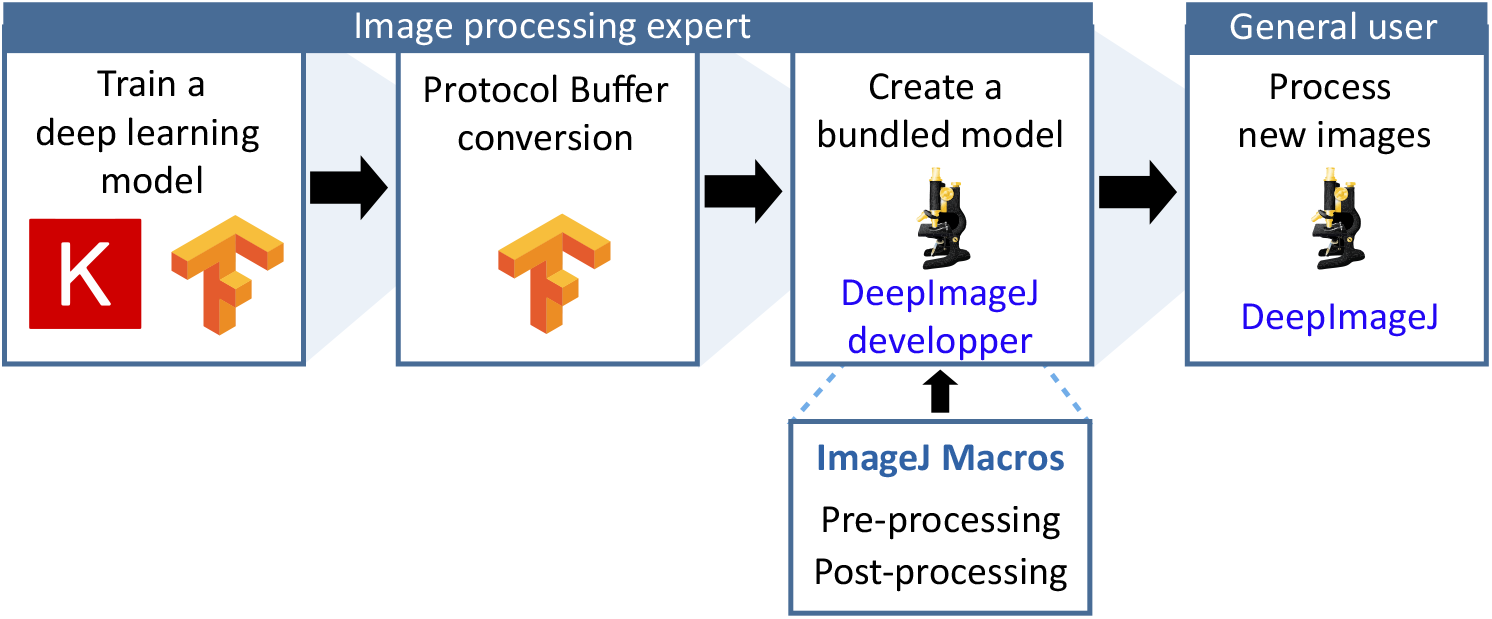
Proposed pipeline for image analysis using deep-learning models and deepImageJ plugin.

The data used was made available by the Cell Tracking Challenge initiative (CTC)^12^ [8, 9]. They are 2*D* phase contrast microscopy videos of Glioblastoma-astrocytoma U373 cells grown on a polyacrylamide substrate. We refer to the model trained on the U373 cells as `U-Net glioblastoma segmentation`. Only a small portion of the data was chosen to train and test the model (see Table 2 for details). Moreover, the image size was halved to shorten the computational time during training. The `Keras ImageDataGenerator` class was used to perform data augmentation with random rotations of *±*40^*◦*^, shifts, shear deformations, zooms, vertical and horizontal flips.

**Table 2:**
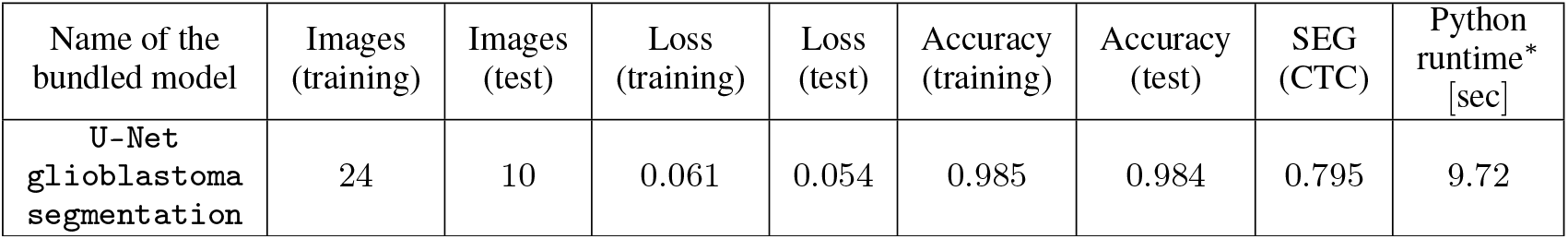
Summary of U-Net training. ^*^Run on an Intel(R) Core(TM) i7-4790 CPU @ 3.60GHz, 32.0GB (RAM), 64-bit Operating System, x64 based processor machine with Windows 10 operating system.

Model was trained with the binary cross-entropy loss function, a learning rate of 1*e*^−04^ and a weight decay of 5*e*^−07^ during 10 epochs of 500 steps each. Altogether, it took 17 minutes to train it. The model finally chosen was the one that resulted in the lowest validation loss during training. The probability output maps from the inference were thresholded at 0.5 to get the final binary masks. The segmentation accuracy was assessed using the percentage of correct pixel assignments (accuracy) and the Jaccard index as computed by the code provided at the CTC web page (SEG) [8, 9]. Table 2 summarizes the segmentation accuracy results. A short script along with the code translates the `U-Net glioblastoma segmentation` trained model from the widely used Keras format HDF5 (.hdf5) to the TensorFlow SavedModel one. Note that the script can be easily adapted to translate other models given in Keras format. Then, the generated model can be easily converted to deepImageJ bundled model using the provided builder module (DeepImageJ Build Bundled Model). Subsequently, the model can be loaded in ImageJ and applied to the rest of the images to obtai uniquely labelled celss that can be further processed using the ImageJ ROI Analyzer.

## Supplementary Information

In this section, we include the User Manual of the four modules delivered within the deepImageJ plugin. For simplicity we refer to ImageJ and Fiji, indinstinctly, as ImageJ.

## 5 Install deepImageJ

The deepImageJ installation can be made in one step (Linux and MacOS) or in two steps (Windows).

The first step is common to all OS: installation of deepImageJ in ImageJ. It can be installed automatically in ImageJ2 using the ImageJ update site, or manually. Once you have finished the installation, do not forget to restart your ImageJ session and gcreate a folder called models inside the path of ImageJ (“…/ImageJ/models/”).

The second step in Windows is required to use PyTorch models.

### 5.1 Manual installation

Download the latest release of deepImageJ from the deepImageJ’s web site^13^. You will need to download two packages: `DeepImageJ`*X*.*X*.*X*.*jarandthe*`Dependencies.zip`.*U nziptheZIPfile*.*Copythepluginexecutable* Otherwise, you can directly copypaste all the dependencies in the locations specified above. The last step might produce some version conflicts with existing libraries in your local installation. Thus being careful is advised. If there are already other versions of the dependencies inside the jars folder, some conflicts might appear when ImageJ is started, and the plugin might not function correctly.

### 5.2 ImageJ update sites

Please, add the update site URL (https://sites.imagej.net/DeepImageJ/) manually:

1. In Fiji/ImageJ, click on “Help” > “Update…”.
2. Once the ImageJ Updater pops up, click on “Manage update sites” > “Add update site”.
3. There you can write the name (“DeepImageJ”) and the URL given (https://sites.imagej.net/DeepImageJ/).
4. Restart Fiji/ImageJ to see the new plugin: `ImageJ > Plugins > DeepImageJ`.

### 5.3 Windows installation

After the installation of deepImageJ in ImageJ is concluded, the Visual Studio 2019 redistributable needs to be installed. You can go to their main site^14^, download Visual Studio 2019 redistributables and install them following their guidelines.

## 6 The deepImageJ environment in ImageJ

There are four different options to work with deepImageJ:

- Run: Runs a specific model to process an image already opened in ImageJ.
- Build BundledModel: User interface to convert TensorFlow or PyTorch models into deepImageJ bundled models compatible with the Bioimage Model Zoo^15^.
- Install Model: Allows the direct installation of deepImageJ compatible models at the Bioimage Model Zoo.
- Validate: Allows the quantitative comparison between two images using standard accuracy measures.

All deepImageJ plugins look for the folder called models inside ImageJ’s directory and load only those models that follow the deepImageJ’s bundled model format. Example models can be downloaded from deepImageJ’s repository or the Bioimage Model Zoo as a zip file. If the model installation is done manually, you will need to unzip the model file and move the inner folder to models. See Figure 3.

**Figure 3:**
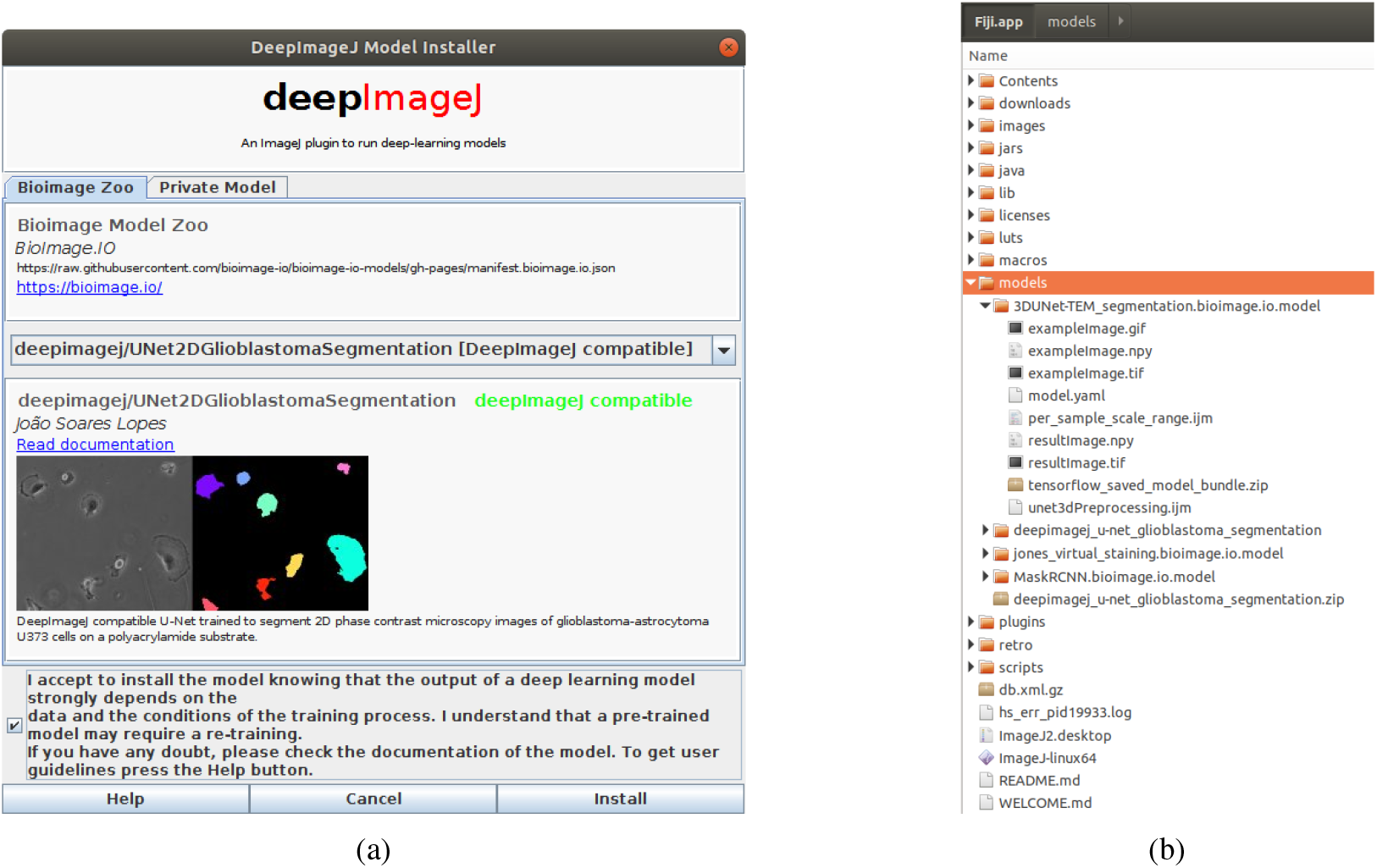
Installation steps: (a) Open ImageJ and use DeepImageJ Install Model to install an example model of the Bioimage Model Zoo public respository. The installation plugin also allows to provide a private url to a model zip. (b) In the Fiji/ImageJ directory, a new folder called models has been created and the model installed is placed there. The plugin automatically unzips the downloaded model so it is ready out of the box to be used by deepImageJ.

### 6.1 DeepImageJ Run

The steps to make inference with a model over an image is as follows:

1. Open the image to process.
2. Click on `ImageJ > DeepImageJ > DeepImageJ Run`.
3. Choose the model to use.
4. Update the tile size if needed. If you do not have the means to know these values, use the values given by default. See the Online Methods for technical details.
5. Click ‘Ok’.

The running window displayed during inference shows in real time the following information: A chronometer, the number of already processed patches, the current memory usage, the memory allocated in ImageJ. If the allocated memory is smaller than the peak memory (see Online Methods), it will not be possible to run the model. This issue can be solved by increasing the allocated memory in ImageJ^16^. Once the process has finished, the input and output images are available for the user to further manipulate them or just to store the result. See Figure 4. The deepImageJ Run functionality is macro recordable.

**Figure 4:**
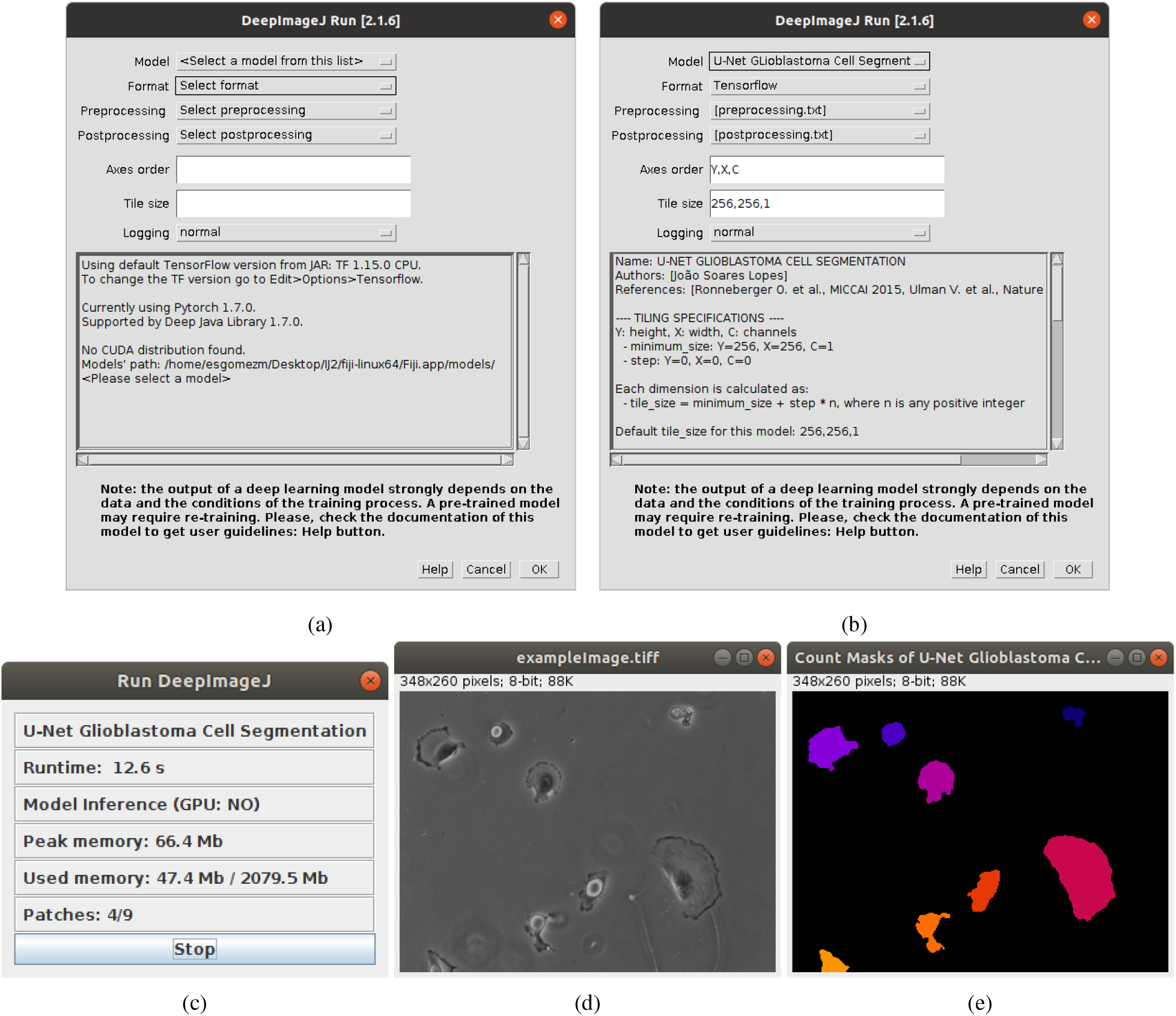
DeepImageJ Run: (a) With an image opened in ImageJ, launch deepImageJ Run. (a) A window indicating the installed TensorFlow, PyTorch and CUDA version appear. (b) Choose a model to run. (c) While deepImageJ processes the image, the progress window informs the user about the runtime, whether a local GPU connection was established, the peak of memory in time and the maximum one needed, and the progress in the patches (tiles) processing. When the process finishes, the user can see (d) the input image and (e) the resulting one.

### 6.2 DeepImageJ Build Bundled Models

The developer module of deepImageJ has a user-friendly interface in Fiji/ImageJ that allows to convert TensorFlow and PyTorch models into deepImageJ compatible format (bundled models) (see Figure 5). The interface asks for a directory where our model is stored. If it is a TensorFlow model saved as saved bundled, the directory should contain the variables sub-folder and a `saved_model.pb` file. If the model is in TorchScript format, then the directory should contain a file of the form `my_trained_model.pt`. The output of deepImageJ Build Bundled Model once all the steps are executed, is the bundled model described in the online methods.

**Figure 5:**
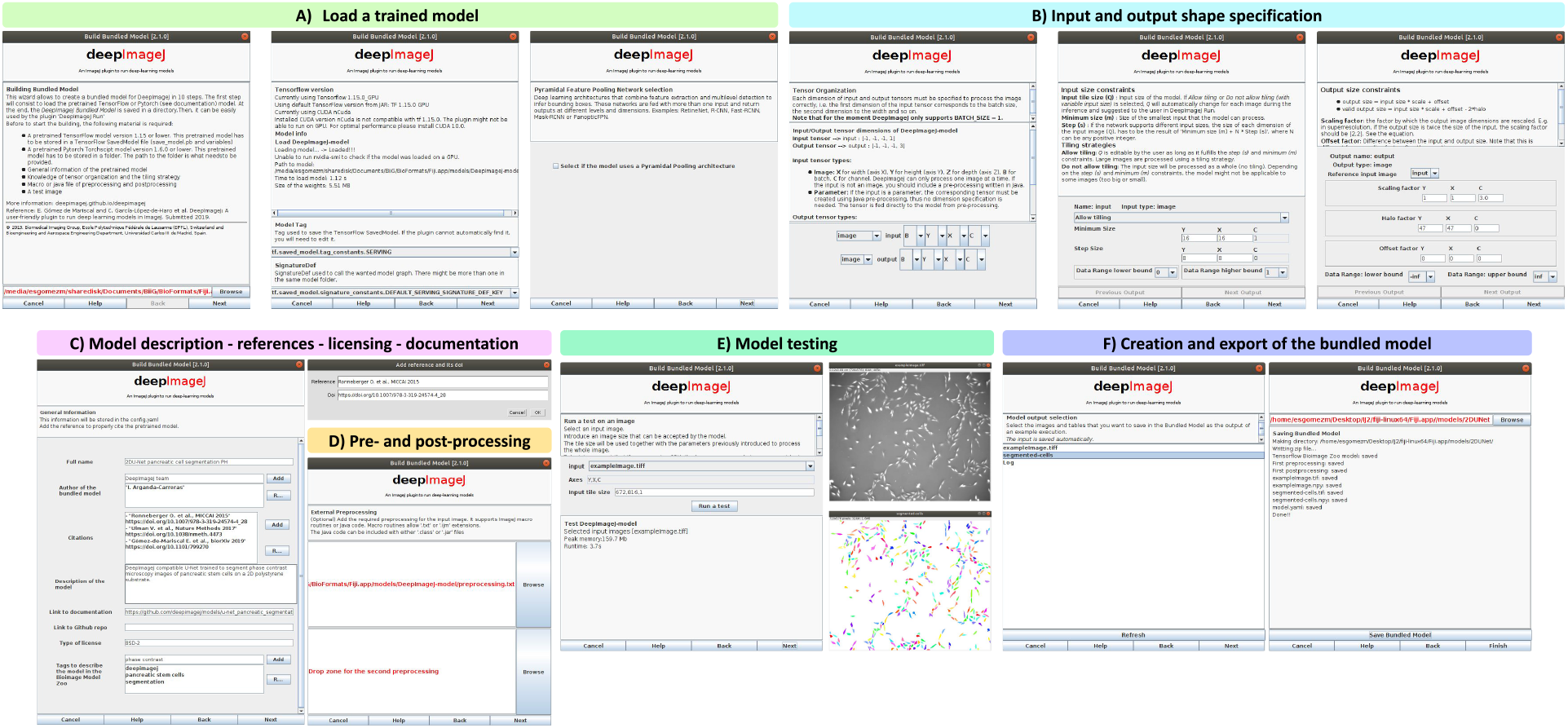
A) The developer module of deepImageJ loads a model from the specified path to the SavedModel directory (variables folder and saved_model.pb file). If it is a TensorFlow model, it asks for TensorFlow session tag constant name. If it is a PyTorch model, it will ask for the number of inputs and outputs that the model has. B) Input and output dimension order (N: batch number, Y: height, X: width, C: channels), input and output shapes definition is specified. C) The model needs to be properly documented for reproducibility and licensing purposes. This step is particularly important for those models that will be publicly available in the Bioimage Model Zoo. D) The developer may also include optional pre- and post-processing macro routines. E) A sample image must be provided, so deepImageJ can test the model. F) If the output is correct, the model is ready to be exported as a deepImageJ bundled model.

### 6.3 DeepImageJ Validate

The plugin will compare quantitatively two images. It asks for a reference and a test image. It is recommended to let the ground truth or the annotated image as a the reference image. The quantitative measures available will either evaluate a discrete result (segmentation) or a continuous one (regression). We included most of the loss functions available in TensorFlow Java API and some common ones such as the signal-to-noise (SNR) ratio. See Figure 6.

**Figure 6:**
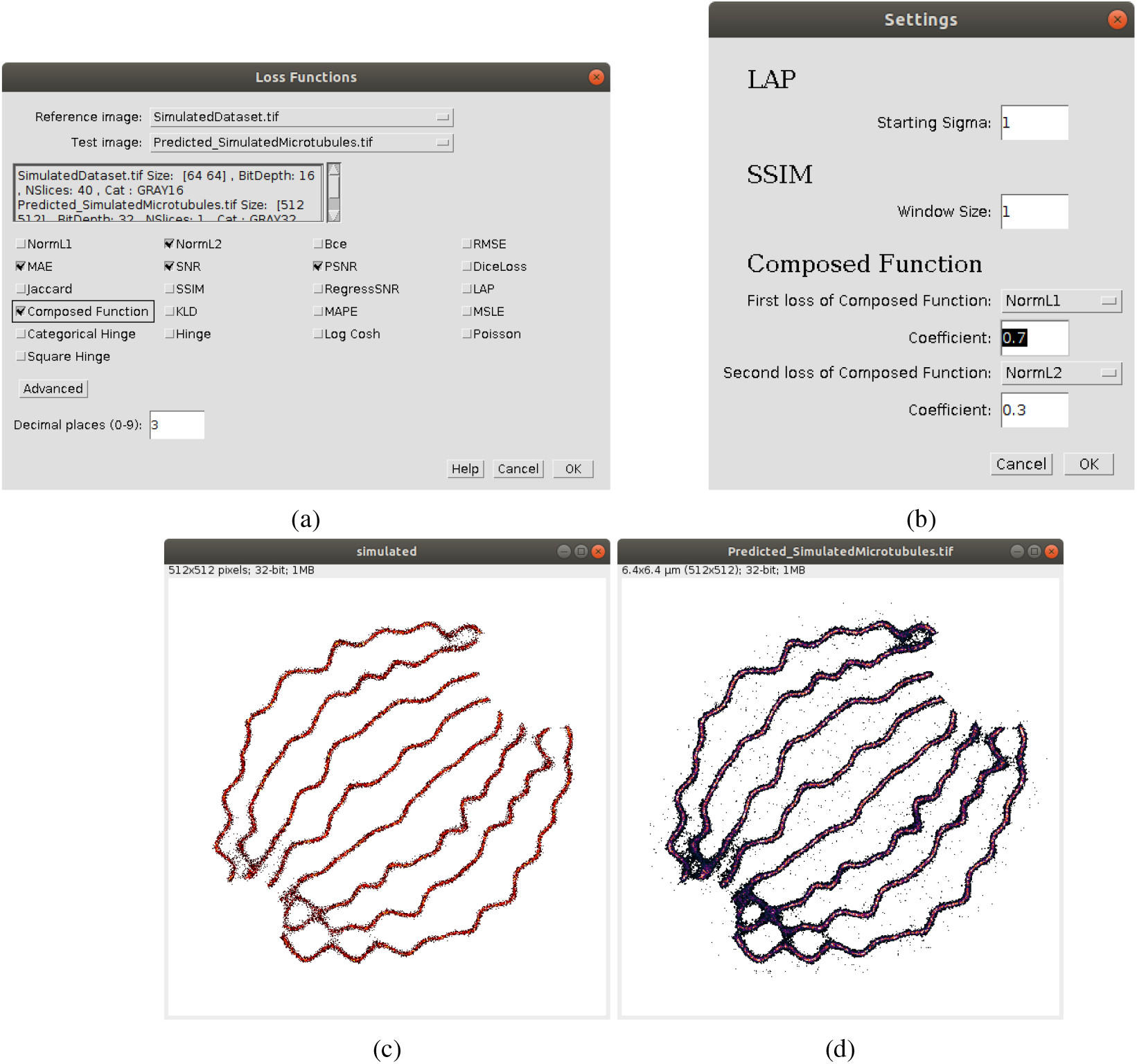
DeepImageJ Validate. (a) The plugin asks for a reference image (c) and a test image (d). (b) The plugin allows for a weighted sum of functions.

## Footnotes

1 https://csbdeep.bioimagecomputing.com/

2 https://org.ee.ucla.edu/

3 https://github.com/adines/DeepClas4Bio

4 https://www.ilastik.org/

5 https://imjoy.io/

6 https://github.com/HenriquesLab/ZeroCostDL4Mic/wiki

7 http://icy.bioimageanalysis.org/

8 https://colab.research.google.com/

9 https://deepimagej.github.io/deepimagej/

10 https://www.tensorflow.org/

11 https://pytorch.org/

14 http://bioimage.io/

15 Building a BioimageAnalysis Workflow using Deep Learning: https://github.com/NEUBIAS/neubias-springer-book-2021/tree/master/Ch03_Building_a_Bioimage_Analysis_Workflow_using_Deep_Learning

16 https://github.com/deepimagej/pydeepimagej

17 Building a BioimageAnalysis Workflow using Deep Learning: https://github.com/NEUBIAS/neubias-springer-book-2021/tree/master/Ch03_Building_a_Bioimage_Analysis_Workflow_using_Deep_Learning

1 https://deepimagej.github.io/deepimagej/

2 https://github.com/deepimagej/deepimagej-plugin/wiki

3 https://github.com/imagej/imagej-tensorflow

4 https://djl.ai/

5 https://visualstudio.microsoft.com/

6 https://github.com/deepimagej/python4deepimagej/

7 https://github.com/bioimage-io/configuration

9 https://github.com/HenriquesLab/ZeroCostDL4Mic/wiki

10 https://github.com/zhixuhao/unet

11 https://github.com/deepimagej/python4deepimagej/

12 http://celltrackingchallenge.net/

13 https://deepimagej.github.io/deepimagej/

14 https://support.microsoft.com/en-us/help/2977003/the-latest-supported-visual-c-downloads

15 https://bioimage.io

16 https://imagejdocu.tudor.lu/faq/technical/how_do_i_increase_the_memory_in_imagej

